# Is *HSPG2* a modifier gene for Marfan syndrome?

**DOI:** 10.1101/849133

**Authors:** Isabela Gerdes Gyuricza, Rodrigo Barbosa de Souza, Luis Ernesto Farinha-Arcieri, Gustavo Ribeiro Fernandes, Lygia V. Pereira

**Author notes:** Correspondence to LVP.

## Abstract

Marfan syndrome (MFS) is a connective tissue disease caused by mutations in the *FBN1* gene. Nevertheless, other genes influence the manifestations of the disease, characterized by high clinical variability even within families. We mapped modifier loci for cardiovascular and skeletal manifestations in the mgΔ^loxPneo^ mouse model for MFS and the synthenic loci in the human genome. Corroborating our findings, one of those loci was identified also as a modifier locus in MFS patients. Here we investigate the *HSPG2* gene, located in this region, as a candidate modifier gene for MFS. We show a correlation between *Fbn1* and *Hspg2* expression in spinal column and aorta in non-isogenic mgΔ^loxPneo^ mice. Moreover, we show that mice with severe phenotypes present lower expression of *Hspg2* than those mildly affected. Thus, we propose that *HSPG2* is a strong candidate modifier gene for MFS and its role in modulating disease severity should be investigated in patients.

## INTRODUCTION

Marfan Sydrome (MFS – MIM# 154700) is an autossomal dominant disorder of the connective tissue with high clinical variability both between and within families (1). It is caused by mutations in *FBN1* gene encoding fibrilin-1, the major component of microfibrils (2). Microfibrils are present in several tissues, which makes MFS a pleiotropic disease affecting mostly the ocular, cardiovascular and musculoskeletal systems (3).

Previous works had suggested that variations on *FBN1* expression caused by polymorphisms in the gene could play a role as a modifier of disease severity (4,5). However, giving the poor genotype-phenotype correlations and the large intrafamilial clinical variability of the disease, recent works have been focusing on understanding how variants in other genes influence MFS phenotypes (6–8).

The effect of genetic background on phenotypic variability in MFS was demonstrated in mice for the first time by our group (9). We showed that mgΔ^loxPneo^ mice in the 129/Sv (129) isogenic background presented earlier age of onset of the disease when compared to those in the C57BL/6 (B6) background. Subsequently, we identified *loci* modulating the phenotypic variability in B6/129 mixed mgΔ^loxPneo^ mice. One *locus* on chromosome 4 is associated with variability of the cardiovascular phenotype, while another, on chromosome 3 is associated with skeletal variability (8). Interestingly, these *loci* are synthenic with two contiguous regions on human chromosome 1 (8).

One of the candidate genes we identified in this region was *Hspg2*, which encodes perlecan, a heparan-sulfate proteoglycan. Mutations in *Hspg2* are associated with Schwartz-Jampel Syndrome (SJS) (SJS1; MIM# 255800), an autosomal recessive disease characterized by skeletal manifestations. Knockout mice for *Hspg2* present severe kyphoscoliosis and dwarfism (10). In addition, these mice die around birth due to severe heart arrest, showing that *Hspg2* plays a role in the formation of not only the skeletal but also the cardiac system (10).

Biochemical studies show that perlecan is also involved in maintenance of vascular homeostasis by its interaction with several extracellular matrix (ECM) components, including fibrillin-1 (11,12). This interaction is essential for the positioning of fibrilin-1 multimeres in the pericellular space and, consequently, the assembly of microfibrils (12–14).

Given these findings, we propose that *Hspg2* is a strong candidate modifier gene for MFS. Here we used the MFS mouse model mgΔ^loxPneo^ on a mixed background (B6/129) to compare *Hspg2* expression between mild and severely affected mice. We show a correlation between *Fbn1* and *Hspg2* expression, and an association between lower *Hspg2* expression and more severe vascular and skeletal phenotypes, corroborating our hypothesis of *Hspg2* as a modifier gene of MFS.

## MATERIAL AND METHODS

### B6/129 mgΔ^loxPneo^mice tissues collection and phenotyping

B6/129 mgΔ^loxPneo^ mice were generated as previously described (8). The project was approved by the Ethics Committee for Animal Experimentation of the Institute of Biosciences, University of São Paulo.

Skeletal phenotype: Full body digital radiographic images of euthanatized animals were obtained using In-Vivo Imaging System FX PRO (Bruker, Germany) for skeletal phenotyping. Kyphosis index (KI) was measured as described (15). We selected 10 mice with the lowest KI values as the severe group and the 10 mice with the highest KI values as the mild group. Thoracic spinal column fragments were collected and frozen in liquid nitrogen for RNA extraction.

Vascular phenotype: Thoracic aorta and thoracic spinal column fragments were collected from each mouse. One fragment was frozen in liquid nitrogen for RNA extraction, while another was processed as described (16) for histological analysis. For each animal, three transversal slices of the aorta were analyzed for elastic fibers fragmentation by optical microscopy, where number of fragmentations (N) was counted (Supplemental Figure 1). Elastic fibers integrity index (EFI) was calculated as following: 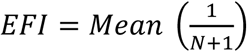.

**Figure 1.**
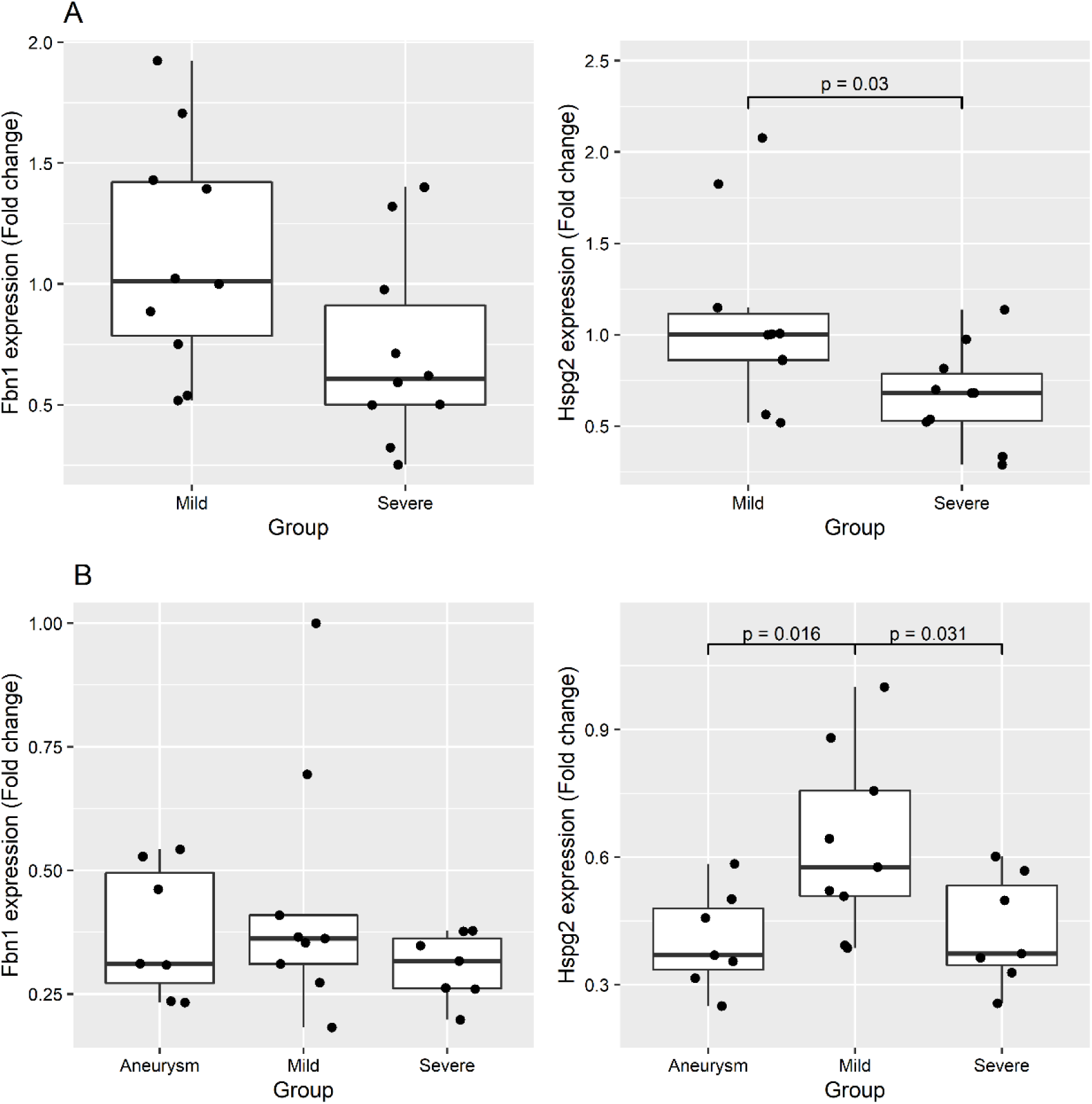
Expression of *Hspg2* and *Fbn1* in mgΔ^loxPneo^ mice. Box plots with expression of *Fbn1* (left) and *Hspg2* (right) in animals with mild and severe phenotypes in (A) spinal column; and (B) aorta. Aneurysm: mgΔ^loxPneo^_A group.

Thoracic aortic transversal sections were also screened for aneurysms. Mice with aneurysms (n = 7) were considered as a separated group (mgΔ^loxPneo^_A). Mice presenting an EFI higher than the third quartile of distribution were considered mild (n = 9) and mice presenting an EFI lower than the first quartile of distribution were considered severe (n = 9).

### RNA extraction and gene expression assay

Spinal column and thoracic aorta fragments were macerated in liquid nitrogen in Trizol® reagent (ThermoFisher) and the RNA was isolated by RNeasy MiniSpin using RNeasy MiniKit (Qiagen). Complementar DNA (cDNA) was obtained from 500ng of total RNA using SuperScript™ III Reverse Transcriptase (Thermofisher). For both *Fbn1* and *Hspg2* expression analysis we used TaqMan® Gene Expression Assay (Thermofisher - Mm01334119_m1 and Mm01181173_g1, respectively). The gene *Actb* (TaqMan® Gene Expression Assay - Mm00607939_s1) was used as endogenous control.

### Statistical analysis

Wilcoxon non-parametric test was used for statistical analysis on comparative gene expression. Pearson correlation test was also used for gene expression correlation analysis. Tests which obtained p-value<0.05 were considered statistically significant and rejected null hypothesis. All the computation was performed using R software (version 3.6.1).

## RESULTS

### Hspg2 expression and its association with phenotypes severity

Heterozygous mgΔ^loxPneo^ (F2 B6/129) animals were phenotyped for skeletal and cardiovascular systems and separated in groups according to severity (Supplemental Figure 1). Expression of *Fbn1* and *Hspg2* was quantified in aorta and spinal column fragments (Figure 1). No difference of *Fbn1* expression in the spinal column was observed between severe and mild mice for skeletal phenotype (Figure 1A). In contrast, we observed lower expression of *Hspg2* in mice with more severe hyperkyphosis in comparison with the mild group (p < 0.05) (Figure 1A).

*Fbn1* expression in the aorta did not vary among the three groups of mgΔ^loxPneo^ mice separated according to vascular phenotype (mild, severe and with aneurysm) (Figure 1B). However, expression of *Hspg2* was lower in the severely affected group and the group with aneurysm when compared to mildly affected animals (p < 0.05) (Figure 1B).

### Correlation between Fbn1 and Hspg2 expression

Although we did not find any differences in *Fbn1* expression between groups in the two phenotypes analyzed, we tested for correlation between *Fbn1* and *Hspg2* expression which could suggest co-function of the corresponding proteins in tissue maintenance. We observed a significant positive correlation between *Fbn1* and *Hspg2* expression on both spinal column and aortic fragments (p < 0.01 | r^2^ = 0.66 and p < 0.01 | r^2^ = 0.91, respectively) (Figure 2).

**Figure 2.**
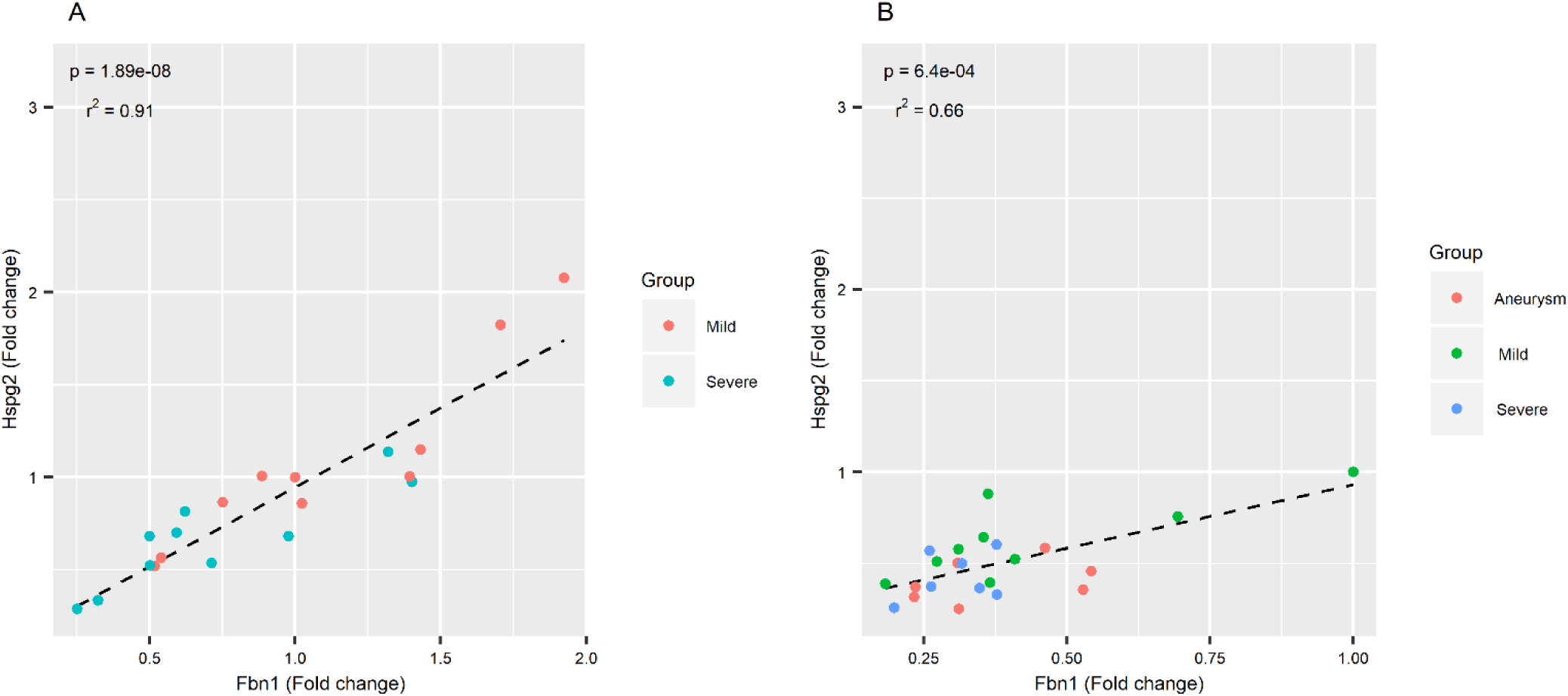
Correlation between *Fbn1* and *Hspg2* expression in mgΔ^loxPneo^ mice. Scatter plots showing positive correlation between *Fbn1* and *Hspg2* expression in (A) spinal column from mice with severe and mild skeletal phenotype (r^2^ = 0.91 | p = 1.89×10^−8^); and (B) in the aortic tissue from the three groups (mild, severe and aneurysm) for the vascular phenotype (r^2^ = 0.66 | p = 6.4×10^−4^). Aneurysm: mgΔ^loxPneo^_A group.

## DISCUSSION

The development of the mgΔ^loxPneo^ mouse model of the intrafamilial clinical variability of MFS allowed us to identify the Awtq1 locus in mouse chromosome 4/human chromosome 1 as a modifier of the cardiovascular phenotype. Within that locus we highlighted *Hspg2* as a candidate modifier gene based on its involvement in skeletal and cardiovascular function, and its direct interaction with fibrillin-1 (10,12,17).

More recently, a combination of genome-wide approaches in 1070 MFS patients led to the identification of a smaller modifier locus for the cardiovascular phenotype named gMod-M1 which overlaps with Awtq1 (7). The authors reported that the only candidate gene within that locus was *ECE1*, highly expressed in the aortic wall and putatively involved in regulation of endothelial-to-mesenchymal transition (7). However, the region also contains the perlecan enconding *HSPG2* gene which, interestingly, has a 2.5-fold higher expression in aorta than *ECE1* (227.2 TPM vs. 93.88 TPM, n=432; GTEx Portal).

Our expression data from mgΔ^loxPneo^ mice with varying aortic and skeletal phenotype severity show a positive correlation between *Hspg2* and *Fbn1*, strengthening the hypothesis of co-function of the corresponding proteins in those systems (12–14,18). Moreover, although we did not find any correlation between *Fbn1* expression levels and phenotype severity, we showed that lower expression of *Hspg2* is associated with more severe hyperkyphosis and with low integrity of elastic fibers and presence of aneurysms. Thus, our findings indicate that expression of *Hspg2* may influence the severity of both skeletal and vascular phenotypes in MFS mice.

The identification of modifier genes of monogenic diseases not only gives clues about the molecular mechanism of pathogenesis and novel therapeutic strategies, but also contributes to the prediction of disease severity. Here we build a case for *HSPG2* as a modifier gene for MFS by reviewing the literature and presenting corroborating evidence in our mouse model of MFS intrafamilial clinical variability. We propose that the role of *HSPG2* in modulating the severity of skeletal and cardiovascular manifestations should be investigated in large cohorts of MFS patients.

## SUPPLEMENTAL FIGURE LEGEND

**Supplemental Figure 1. Quantification of skeletal and vascular phenotypes.** (A) X-ray showing lines used for the calculation of KI (KI=|AB|/|CD|); (B) histology of cross sections of the aorta in mgΔ^loxPneo^ animals. Red arrow points to aneurysm; red arrowheads show sites of elastic fiber fragmentation. Scale bars 50μm (above) and 10μm (below).

