## Supplementary figures and images for "Is *HSPG2* a modifier gene for Marfan syndrome?"

### Supplemental Figure 1

#### Slide 1
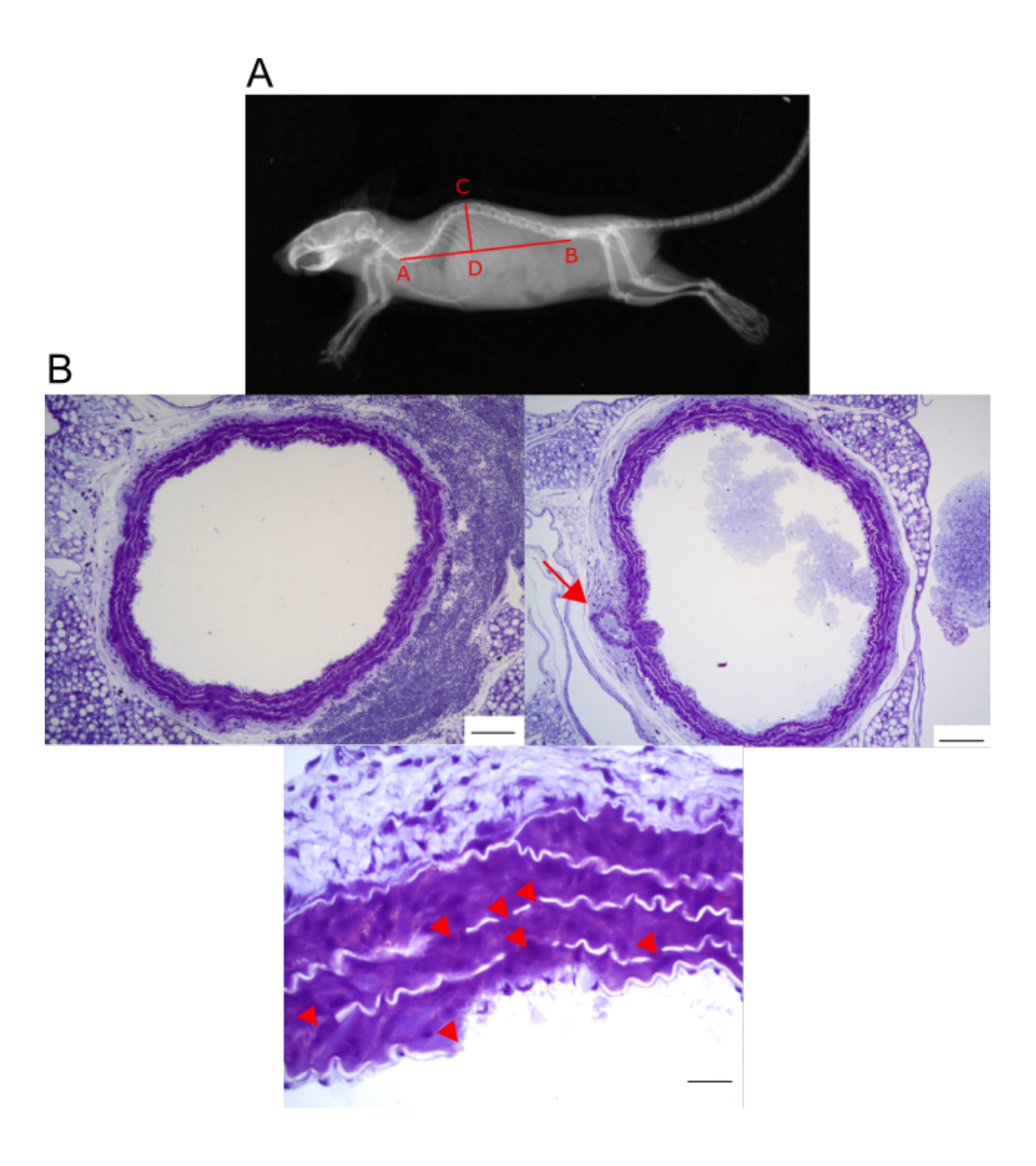
